# Deep behavioral phenotyping tracks functional recovery following tibia fracture in mice

**DOI:** 10.1101/2025.05.09.652892

**Authors:** Jonathan E. Layne, Dustin M. Snapper, Molly E. Czachor, Charles Lam, Jacob D. Matityahu, Dane R.G. Lind, Matthieu Huard, Kazuhito Morioka, Julian C. Motzkin, Allan I. Basbaum, Jarret A.P. Weinrich, Chelsea S. Bahney

## Abstract

**Introduction:** An estimated 178 million fractures occur worldwide each year, with lower limb fractures in particular showing a high incidence of poor healing, and these often lead to reduced mobility and chronic pain. Bone healing and the ability to bear weight are closely tied to the mechanical stability of the fracture site. Although fracture stabilization is a well-established factor modulating the rate and extent of bone repair, there is a notable gap in non-destructive technologies that can rapidly and objectively quantify functional recovery in preclinical settings. We consider this to be a significant limiting factor in translational studies directed at improving fracture healing. Here, we describe a novel behavioral phenotyping approach that enables rapid quantification of post-fracture weightbearing and kinematic metrics in freely behaving mice. Our goal is to identify and characterize metrics most indicative of fracture-induced behavioral impairment and to use these metrics to quantify how functional recovery is altered in mice with pin stabilized versus non-stabilized fractures. We also use this approach to explore whether sex is a significant contributor to functional recovery.

**Methods:** Male and female adult C57BL6/J received a mid-shaft tibial fracture that was either left unstabilized or fixed with an intramedullary pin. Non-fractured naïve mice served as controls. Behavioral recordings of freely moving mice were acquired prior to fracture and then throughout the time course of healing, from 5 to 35 days post fracture (DPF). To track mice and analyze changes in paw pressure and kinematic behaviors after fracture, we then applied a novel machine learning-enabled behavioral phenotyping analysis.

**Results:** In this study, we demonstrate that severity of the behavioral phenotype is more significant in mice with unstabilized fractures when compared to mice with pin-stabilized fractures. Pin stabilization generally allowed increased weightbearing and produced smaller changes in kinematic metrics. Interestingly, we observed only minor sex specific differences in fracture-induced behavioral impairments and recovery. Our analysis also revealed that functional recovery is more complex than is a set of individual parameters viewed in isolation. In fact, unique behavioral parameters identified different time windows for functional recovery. Therefore, we developed a comprehensive, unified graph theoretic metric of fracture recovery that encompasses all behavioral parameters quantified. Using this unified metric, we confirmed the increased severity of the fracture phenotype in unstabilized versus pin stabilized mice and identified a clear time window of functional recovery, for both fracture groups.

**Discussion:** Our findings demonstrate how this novel comprehensive behavioral phenotyping approach, which combines machine learning and graph theory, makes it possible to rapidly quantify longitudinal changes in mice after fracture. This approach enables us to determine functional recovery patterns based on a unified behavioral metric of healing. Our data and methodology form a foundation for future mechanistic experiments focused on understanding biological or mechanical variables that influence functional healing and will also enable more rapid testing of various strategies to accelerate bone healing.

## INTRODUCTION

Bone fractures are among the most common orthopedic injuries, with lower limb fractures accounting for 47.3 million fractures globally in 2019.[1] Delayed healing, or failure to heal, is especially common in lower limb fractures with 2-year complications reported to occur in 13.6% of femur and 11.7% of tibia fractures.[2] Many non-modifiable factors affect the rate of bone union, such as fracture pattern, degree of soft tissue damage, age, sex, smoking status, and medical comorbidities. [3] However, the orthopaedic surgeon has the greatest control over the mechanical environment of the fracture site, through implant choice, insertion method, and timing of post-operative weightbearing. Mechanical loading is essential for effective bone healing and reduces the risk of delayed union or nonunion following a lower-limb fracture.[4] As a result, patients are encouraged to bear weight, including both partial and full weight-bearing as tolerated. However, increased pain levels have been shown to lead to decreased weight-bearing and less effective participation in physical therapy, which can negatively impact recovery.[5, 6]

Long bone fractures heal through four distinct but overlapping biological phases as we recently reviewed. [7, 8] Briefly, following fracture, a hematoma forms to stop the bleeding, contain bone fragments, and trigger a pro-inflammatory cascade that is critical to initiating the repair response.[9, 10] In mice, this hematoma/pro-inflammatory phase typically spans the first 5 days post fracture. Bone healing then proceeds through two distinct processes, which occur in parallel as the hematoma begins to resolve. Along the cortical surfaces of the bone, skeletal progenitor cells differentiate to form bone directly through the process of intramembranous ossification.[11] Within the fracture gap, skeletal progenitor cells differentiate into chondrocytes to form a provisional cartilage matrix that bridges the broken bone ends and provide relative stability. Cartilage in the fracture gap then transforms to bone through the process of endochondral ossification.[12–18] In the final phase of healing, osteoclasts remodel the newly formed trabecular bone into cortical bone.[19]

As to preclinical mechanisms, currently there is a poor understanding of how the biological process of fracture healing correlates with functional behaviors in mice. And, importantly, there is a growing body of evidence suggesting that the paucity of preclinical functional outcome measures in fracture repair hinders translation of effective treatments from mice to humans.[20–24] The standard quantifiable preclinical outcome measures of bone healing are rarely based on behavioral assessment, but rather are destructive, including, histological tissue evaluation, *ex vivo* microcomputed tomography (μCT), gene and protein expression analysis, and biomechanical bone quality testing. These analyses are time consuming, expensive, and typically require a large number of animals to obtain the different destructive data sets. Gait analysis, such as through DigiGait or CatWalk, has been used infrequently in fracture healing studies, due to the specialized equipment needs, the challenge in applying these protocols to mice with fracture, and the time-consuming analysis.[25] For this reason, there remains an important technology gap for rapid, unbiased, and non-destructive evaluation of clinically informed outcome measurements that can provide more quantitative assessments of fracture-induced pain and functional recovery.

Here, we present data from a longitudinal behavioral phenotyping study in mice in which we quantitatively track functional recovery (i.e., weightbearing and kinematic shifts) after long bone fractures of the lower limb. Our goal was to assess the impact of mechanical stability and sex on functional behavioral recovery after tibia fracture. To represent the clinically modifiable mechanical environment of the fracture site, we compared two different methods of fracture fixation, namely, intramedullary pin stabilized versus unstabilized fractures. Additionally, we assessed the contribution of the non-modifiable factor of sex, as presently the preclinical and clinical data are conflicting on the role that sex plays in fracture healing. [25–27] Analyzing individual behavioral metrics, based upon fracture fixation in both male and female mice, we find clear and significant differences in the severity and time course of multiple distinct weightbearing and kinematic changes. With regards to sex specific differences, although we did observe differences in weightbearing and kinematic behaviors between male and female mice at baseline, after normalization, differences in fracture-related behavior and recovery across individual metrics were minimal. We then developed a novel graph theoretic approach that integrated the many distinct behavioral measures into a single, comprehensive metric that quantifies the global behavioral state of each mouse during recovery from fracture. Using this unified metric, we confirm the relative importance of fracture stability to functional recovery and rigorously establish the time window for functional recovery after tibia fracture. Taken together, this new behavioral phenotyping methodology provides the foundational basis for unbiased, sensitive, user-independent quantitative assessments of functional impairment and recovery after bone fracture.

## Materials and Methods

### Animal Husbandry and Ethical Approval

Adult (10–14-week-old) male and female wild-type C57BL6/J (Jackson Laboratories Stock #000664) mice were used for all experiments. Experiments complied with ethical regulations and protocols were approved by the Institutional Animal Care and Use Committee (IACUC) at our university. All mice were group-housed, provided environmental enrichment, fed a standard diet, and maintained in facilities with standard light/dark cycle and appropriate environmental controls.

### Surgical Procedures

Mice either received an unstabilized or intramedullary stabilized tibia fracture. Non-fractured mice were used as a control group. Mice were anesthetized prior to fracture using isoflurane inhalation (4-5% induction, 2-3% maintenance). All surgeries were performed on a heated operating table using aseptic technique and ocular ointment was placed on the eyes during anesthesia. Sustained release buprenorphine (3.25 mg/kg, Fidelis Animal Health, Cat#NDC 86084-100-30) was administered through intraperitoneal injection after completion of the surgery. Following surgery, the mice were socially housed and allowed to ambulate freely and after surgery, were monitored for 72 hours for pain and discomfort.

#### Unstabilized tibia fracture

To create a mid-diaphyseal fracture of the right tibia, anesthetized mice were placed pronated under a custom-built three-point bending fracture apparatus. No fixation was provided after the creation of the fractures, which simulates clinical fractures with a high degree of mobility. As previously described, this technique is a well-established method to create a robust endochondral repair. [28]

#### Stabilized tibia fracture

Following anesthesia induction, the right leg was shaved and sterilely prepped using three rounds of 70% alcohol wipes followed by povidone-iodine swab sticks 10%, (Dynarex Corporation, Cat#1202). The knee of the right tibia was placed in flexion and a small skin incision was made slightly proximal to//from? the tibial plateau to the distal tibia. A 23-gauge needle was used to form a pilot-hole at the top of the tibial plateau. A sterilized insect pin was then inserted through the hole spanning from the tibial plateau through the tibial intramedullary space and secured into the distal tibia. Then, a Dremel was used to create a small hole (0.25-0.5 mm) in the mid-diaphysis of the tibia. Pressure was applied to both the proximal and distal ends to generate a full-thickness tibial fracture as previously described.[29, 30] The pin was then trimmed with wire cutters at the tibial plateau. The incision was closed with 5-0 Biosyn Sutures (Covidien, 5687). Bupivacaine hydrochloride (NovaPlus, RL7562) was applied topically for post-operative pain management.

#### Naïve control mice

Age and sex matched wild-type C57BL6/J mice that received no fracture and no anesthesia were used as controls for each individual fracture type. These mice were ordered at the same time as the fracture mice, housed identically, and monitored at the same cadence as their fractured counterparts.

### Behavioral Video Recording

#### Blackbox device

Mice were assessed for weightbearing and kinematic parameters using the Blackbox R4 device (Blackbox Biotech Inc., BB1R-0015), which captures animal pose and paw pressure. Up to four freely moving mice can be monitored at the same time (**Figure 1**).[31] Briefly, the Blackbox device encloses a single, high speed, high spatial resolution near-infrared (NIR) camera. The mice are placed onto the glass surface, on which 4 black acrylic chambers are placed, with only one mouse per chamber during recording (**Figure 1C**). Paw contact force with the glass is transmitted through frustrated total internal reflectance (FTIR). The FTIR light sources are two 850nm NIR LED strips that are aligned perpendicular to two opposite edges of the glass floor. Transillumination (TL), which enables visual identification of the mouse pose within the chamber, is generated by 4 additional NIR LED strips that are located 10cm below the glass floor. The overall frame rate of the camera is set to 90 frames per second (fps). In every other frame, the TL LEDs are turned off, allowing for exclusive imaging of the FTIR signal. This approach produces an effective frame rate of 45 fps (45 TL + 45 FTIR = 90 total fps). The behavioral recording is captured by BlackBox software (Version 0.1.2) onto a Blackbox workstation.

**Figure 1.**
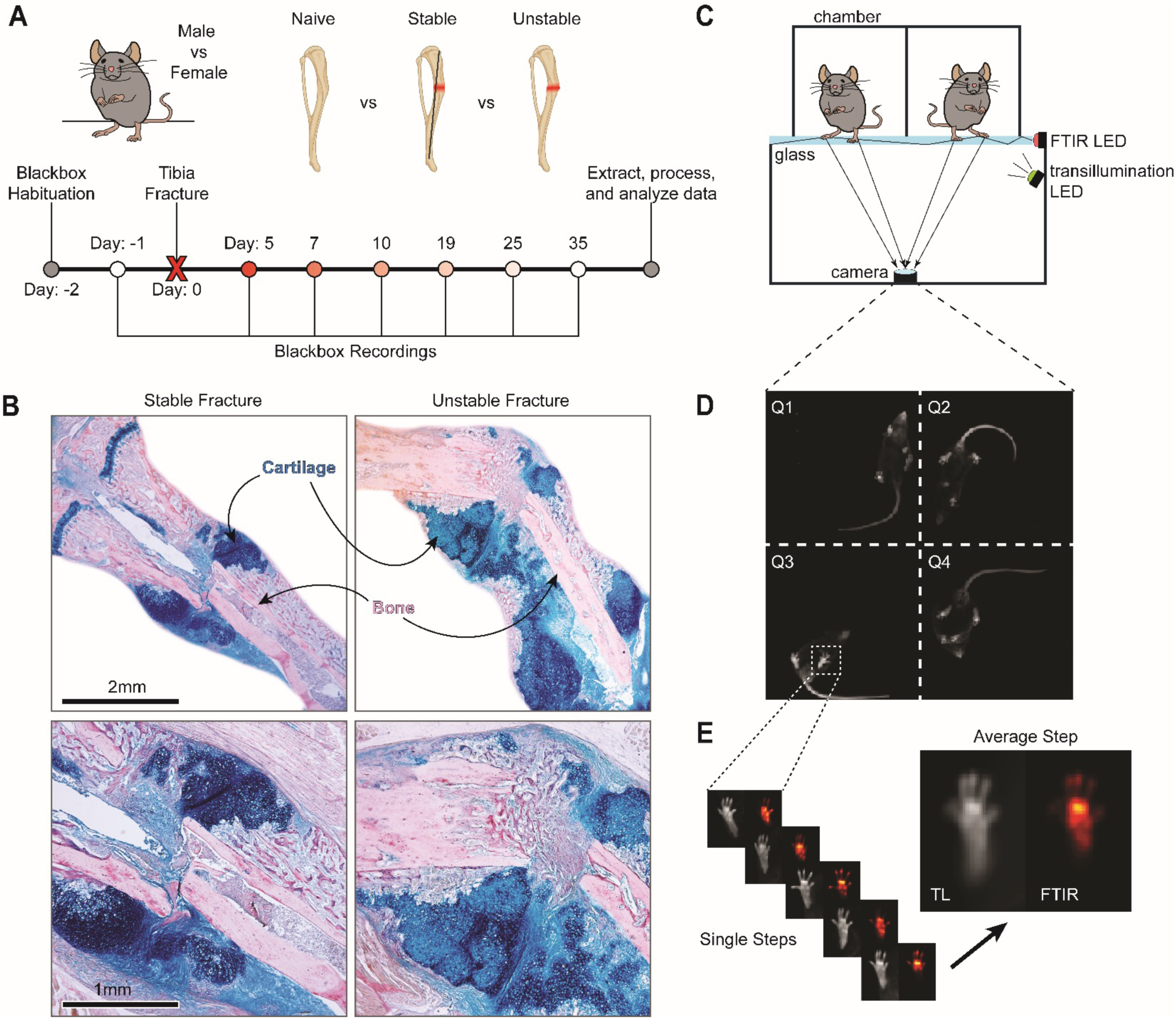
Blackbox monitoring of functional recovery after tibia fracture. **(A)** Schematic representation of experimental groups and Blackbox recording timeline. **(B)** Representative images of Hall-Brundt’s Quadruple (HBQ) stained histological sections of tibiae from female at 10 days post-fracture (DPF) for both stabilized and unstabilized fracture groups. Scale bar = 2mm (top row) and 1mm (bottom row). **(C)** Diagram of the BlackBox device. The device consists of four chambers that house a single mouse during recording, with two of the four chambers pictured here. The glass floor below the chambers allows for the capture of both transillumination (TL) and frustrated total internal reflectance (FTIR) images. **(D)** Representative frame of the 4-quadrant TL video recordings. **(E)** Representative frames of stepping bouts extracted from the TL and FTIR recordings. Both TL and FTIR images were then averaged across steps to generate a representative image of paw placement pressure distribution during stepping.

#### Longitudinal behavioral monitoring

Mice were first habituated to the device for 4-5 minutes for two days before fractures were performed and one day prior to fracture, mice behaviors were recorded (baseline, **Figure 1A**). Post-fracture behavior recording was then completed at 5-, 7-, 10-, 19-, 25-, and 35-days post fracture (DPF). Testing began 5-DPF to ensure the effects of the protocol-required slow-release buprenorphine had worn off. On the day of testing, animals were placed onto the Blackbox, one at a time into an individual chamber (**Figure 1C**). Mice are then recorded continuously for 4-5 minutes. TL and FTIR recordings are saved for further analysis (**Figure 1D-E**).

### Automated Analysis of Behavioral Recordings

To analyze data from Blackbox recordings, we developed a custom-written data processing and analysis pipeline within MATLAB (R2023a, MathWorks) that leverages open-source video processing software (FFMPEG) and machine-learning based video object tracking (DeepLabCut [DLC], v1.5.7). The end result of our pipeline is an analysis of weight-bearing and kinematics of the mouse. See **Table 1** for detailed descriptions of relevant outputs. Data are processed on a Puget Systems Threadripper workstation with an NVIDIA A5000 Ada graphics processing unit (GPU). References to functions below refer to functions native to MATLAB.

**Table 1.** – Definitions of selected behavioral metrics.

| Metric | Description (Please see methods for more details). |
| --- | --- |
| Weightbearing Ratio | The ratio of FTIR light intensity of the hindpaws from the fractured over the non-fractured limb. |
| Pad Intensity | The percentage of FTIR intensity restricted to the pad over the entire paw. |
| Stepping Correlation | The Pearson correlation of stepping speeds during walking for indicated paws (i.e., left vs right hindpaw). |
| Step Duration | The average duration of a step during walking, defined as the full-width half-max from stepping speed. |
| Maximum Speed | The average maximum speed of the paw during stepping during walking. |
| Step Length | Average length of each step during walking. |

#### Video processing and automated object labeling

First, it was necessary to split the single Blackbox videos (both TL and FTIR) containing all 4 chambers into individual videos per mouse, which was accomplished using FFMPEG. Split TL videos are then processed through DLC to identify the pose of the mouse within the chamber. The points identified include: hindpaw (both pad and toes), forepaw (pad and toes), base of the tail, abdomen, chest, and mouth (**Supplemental Figure 1**). For paws, both right and left paws are separately identified. To build the pose estimation model, videos from 24 behavioral recordings were used, including videos from both naïve and fracture mice, with at least 40 still frames per video labeled. Using this model, pose estimation (object tracking) is performed on all behavioral videos.

#### Weightbearing analysis

Weightbearing is measured during the stance phase of walking for each paw. A single weightbearing measurement is taken per step, measured 150 milliseconds after the maximum paw speed during stepping. Weightbearing of an individual paw is measured as the summed intensity of paw luminescence within the FTIR video, with paw placement determined from time-synced TL videos, and the FTIR still frame cropped to fit the measured paw. Within a recording, the summed intensity of weight-bearing is averaged across all identified steps within a recording session. The weightbearing ratio is calculated as the ratio of the mean summed FTIR intensity of compared paws. For example, the hindpaw weightbearing ratio is equal to the ratio of the mean sum of the FTIR intensity of the hindpaw of the fractured limb divided by the hindpaw of the unfractured limb. The weightbearing ratio is calculated for the right vs left hindpaw, right versus left forepaws, right forepaw vs right hindpaw, left forepaw versus left hindpaw, and both forepaws versus both hindpaws.

Additionally, the percentage contribution to weightbearing of the toes, pad, and heel within a single paw is taken. Here, the cropped FTIR image used to determine weightbearing is masked to segregate only the contribution to the overall intensity from the particular part of the paw. This masked intensity value is then summed, divided by the overall intensity of the step, and multiplied by 100% to generate the percent contribution. For the forepaws, only toes and pad contributions are calculated.

#### Kinematic analysis

Next, to extract behavioral endpoints, we analyzed DLC pose for kinematic metrics. Paw-related kinematic metrics extracted are as follows: maximum paw speed during stepping (cm/s), stride duration (full width half max [FWHM] in milliseconds), and stride length (cm). More general locomotor-related measurements include distance traveled (cm, as measured from the movement of the base of the tail), walking speed (cm/s), and percentage of time spent walking during the recording. The final form of the weight-bearing and kinematic metrics is a scalar that represents the average value for each metric within a video (i.e., the average maximum paw speed per paw within a single video).

#### Graph theoretic analysis

First, for each single scalar metric (for example, walking speed), data from all recordings (both sexes; naïve and both fracture types) are z-scored within-metric. This z-scoring is performed separately for all metrics described above. Next, a matrix of pairwise correlation coefficients (Pearson’s rho and correlation significance values) are computed using the z-scored metrics, comparing each recording to all other recordings in the dataset. The significance matrix is corrected for multiple comparisons using the *MAFDR* function, generating a q-value matrix. A positive adjacency matrix is constructed from the correlation matrix by keeping only pairwise correlations greater than 0.3, and q-values less than 0.05. A weighted graph is then constructed using the *graph* function. Within this generated graph, each node corresponds to a single recording, and edges indicate significant positive correlations between nodes, weighted to account for the strength of the correlation. Finally, we use the *distances* function to calculate the distance (the unit of which is total number of weighted edges that make up the shortest path to connect a given pair of recordings) of the baseline recording to post-baseline recordings within a single animal.

### Fracture Histology

Fractured tibia were harvested at 10 DPF, and fixed in 4% paraformaldehyde at 4°C. After 24 hours, tibias were decalcified in 19% Ethylenediaminetetraacetic Acid (EDTA) and left to rock at 4°C for 3 weeks with EDTA changes every other day. Decalcified tibias were dehydrated and then embedded in paraffin. Tissue samples were serially sectioned using a Leica RM 2155 microtome at 8-10 µm, producing microscope slides with 3 sections per slide. Slides were stained using Hall-Brundt’s Quadruple (HBQ) staining protocol callus (**Figure 1B**).[14] Images were captured with a Leica DM5000 B microscope.

### Statistics

Statistical analyses were performed using GraphPad Prism (Version 10.1.2). Data were analyzed for statistical significance using mixed-effect analysis with multiple comparisons and a two-stage linear step-up procedure of Benjamini, Krieger, and Yekutieli. [32] Data are displayed as the mean ± standard error (SEM). The results of all mixed effects models can be found in the tables provided in Supplemental Information (**Supplemental Statistics**). Symbols are used within the graph to indicate statistical significance with p<0.05.

## RESULTS

### Quantifying Functional Recovery Following Fracture in Mice

The aim of this study was to assess functional recovery in mice that received tibia fractures with different stabilization protocols and to determine the contributions of sex to functional recovery. Our experimental groups included male and female mice, both of which are split into groups by fracture type: (1) pin stabilized fractures and (2) unstabilized fractures. Pin stabilized fractures are more translationally relevant as they mimic the intramedullary nail placement standard in human tibia fracture management. As unstable fractures do not include the intramedullary pin, heightened interfragmentary motion about the fracture is possible. Throughout preclinical fracture literature, both stabilized and unstabilized models have been used frequently, however, there are no studies that directly compared functional recovery between these models. Functional recovery after fracture was monitored at 1-day prior to injury (baseline), and then 5-, 7-, 10-, 19-, 25-, and 35-days post fracture (DPF, **Figure 1A**). Biologically, recordings at 5– and 7-DPF represent the soft/cartilage callus stage of healing. Recordings at 10-DPF capture the beginning of the endochondral conversion of cartilage to bone (**Figure 1B**). By 19-DPF, the callus has largely converted to trabeculated bone, with cortical bone remodeling occurring between 25-to 35-DPF.

To monitor behavioral changes, we used the Blackbox system [31] and recorded videos of mice before and after tibia fracture. A single behavioral recording session generates two separate videos, one of which is captured with the transillumination lighting to identify animal pose, and the other captured with the frustrated total internal reflectance (FTIR) lighting, to identify paw contact on the glass (**Figure 1C-E**). To analyze behavioral recordings, we developed a data processing and analysis pipeline (see **Methods** for details).

To understand the presence of sex-specific differences prior to the fracture procedure, we compared gross weights, paw weightbearing, and kinematic parameters of male and female mice at baseline. As has been previously shown, [33, 34] we observe that age-matched male and female mice exhibit significantly different weights and kinematic profiles (**Supplemental Figure 2**). Male mice weighed significantly more than females (28.38 +/− 2.07 grams vs. 22.61 +/− 1.539 grams, **Supplemental Figure 2A**). Similarly, and as would be expected given the difference observed in gross weights, paw luminescence, or the summed FTIR intensity of the whole right hindpaw, was significantly higher in male mice than female mice (**Supplemental Figure 2B**). Additionally, we observed a significant linear relationship between paw intensity and mouse gross weight (**Supplemental Figure 3**). Lastly, we found that males have a shorter step duration and step time, and a slower maximum paw speed than female mice (**Supplemental Figure 2C-E**). Taken together, these data clearly demonstrate that our proposed methodology has the sensitivity to detect significant differences between functionally distinct groups of mice with regards to paw weight-bearing and kinematic parameters.

### Stabilization Decreases the Severity of Functional Deficits and Accelerates Recovery

As noted above, females and males have significant differences in weight-bearing kinematic parameters at baseline (**Supplemental Fig 2**). For this reason, we analyzed the functional recovery from tibia fracture separately in female and male mice. Starting with female mice, over the time course of recovery from fracture, we observed that less weight (i.e., less FTIR signal) is placed on the paw of the fractured limb for both stabilized and unstabilized fracture groups compared to naïve controls (**Figure 2A, Supplemental Videos 1-3).** This decreased weight-bearing on the fractured limb translates to a decreased weight-bearing ratio (ipsilateral fractured limb / contralateral non-fractured limb) for female mice. with both stabilized and unstabilized fracture versus naïve controls (**Figure 2B**). Additionally, we recorded further evidence of changes in the distribution of weight-bearing across the paw of the fractured limb, namely that the percentage of weight-bearing on the pad versus the total paw significantly decreases after fracture (**Figure 2C**). Our kinematic analysis also uncovered significant changes for both fracture types; the fractured limb step duration increases (**Figure 2E**) and maximum paw speed decreases (**Figure 2F**). Furthermore, our kinematic analysis revealed that only in mice with unstable fractures are there significant changes in stepping correlation (**Figure 2D**) and step length (**Figure 2G**). For all weight-bearing and kinematic metrics, the changes induced early in the fracture process recover back to normal over time, albeit at different rates for different metrics. (**Figure 2A-G**)

**Figure 2.**
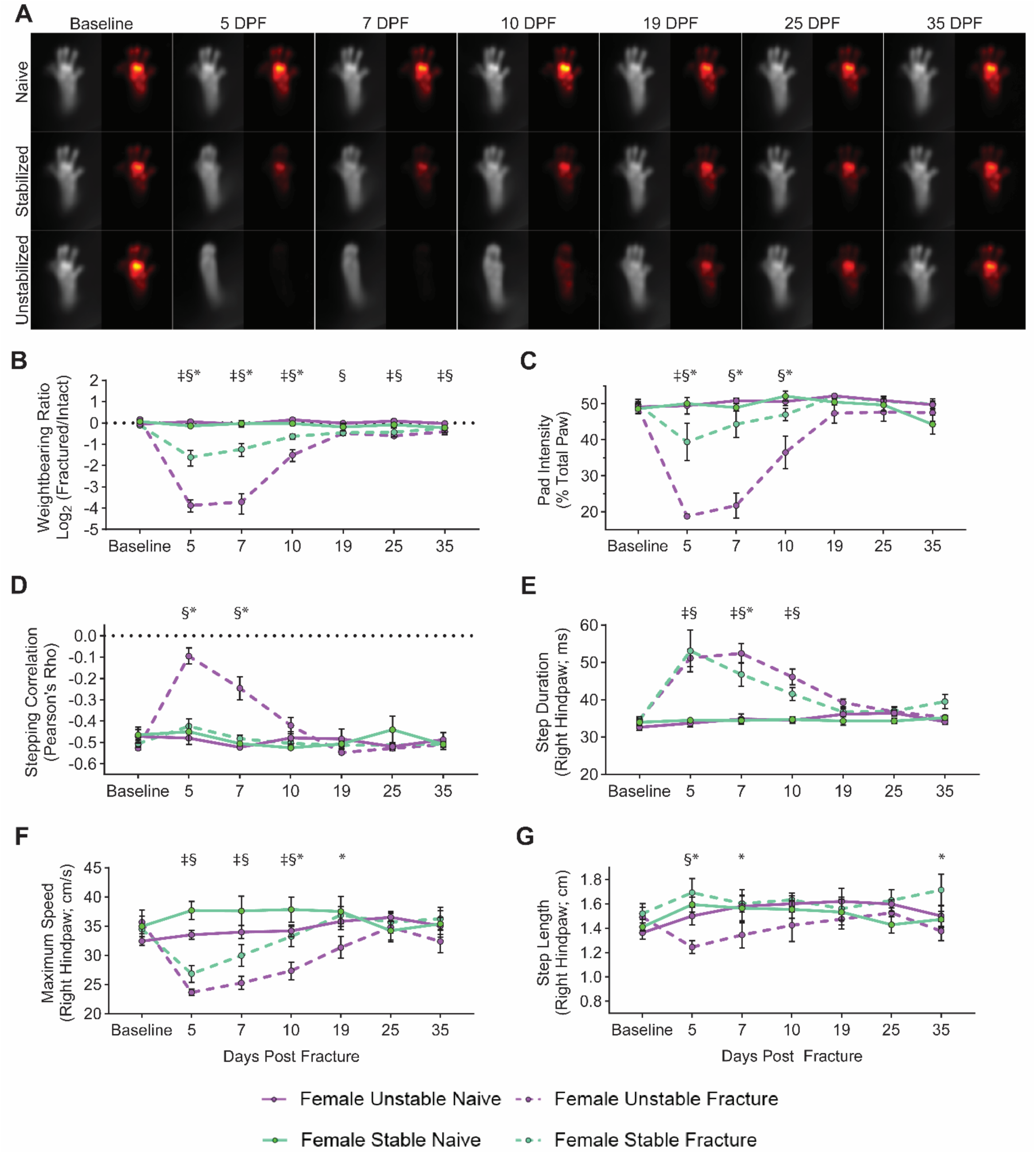
Differential functional impairments in female mice identified by fracture type. **(A)** Average images of paw placement (TL) and weightbearing (FTIR) of naïve, stabilized, and unstabilized fracture females at baseline and 5-, 7-, 10-, 19-, 25-, and 35-days post injury (DPI). Longitudinal analysis of weightbearing and kinematic parameters: **(B)** Weightbearing ratio of the fractured hindlimb to the intact hindlimb (Mixed-effects analysis, Fixed effects: Time (F(6, 48) = 30.89, p<0.0001) Naïve vs. Fracture (F(1, 8) = 80.68, p < 0.0001). **(C)** Percentage of FTIR intensity localized to the pad of the hindpaw of the fractured hindlimb (Mixed-effects analysis, Fixed effects: Time (F(6, 48) = 25.07, p<0.0001) Naïve vs. Fracture (F(1, 8) = 24.65, p = 0.0011). **(D)** Hindlimb stepping correlation (Mixed-effects analysis, Fixed effects: Time (F(6, 48) = 14.13, p < 0.0001) Naïve vs. Fracture (F(1, 8) = 7.037, p = 0.0291). **(E)** The average full-width, half-max duration of a single step, in milliseconds (Mixed-effects analysis, Fixed effects: Time (F(6, 48) = 17.39, p < 0.0001) Naïve vs. Fracture (F(1, 8) = 113.2, p < 0.0001). **(F)** Average maximum speed (cm/s) of right hindpaw during stepping (Mixed-effects analysis, Fixed effects: Time (F(6, 48) = 9.228, p < 0.0001) Naïve vs. Fracture (F(1, 8) = 9.027, p = 0.0170). **(G)** Average distance of each step of the right hindlimb in centimeters (Mixed-effects analysis, Fixed effects: Time (F(6, 48) = 0.9560, p = 0.4648) Naïve vs. Fracture (F(1, 8) = 0.0261, p = 0.9134). Significant differences (p<0.05 after corrections for multiple comparisons) are indicated by: § – unstable fracture and respective naive control; ‡ – stable fracture and respective naive control; * – unstable and stable fracture.

When comparing female mice with stabilized and unstabilized fracture directly, we observed that unstable fracture produces a more severe functional deficit, and that mice with stabilized fracture recover from the fracture event more quickly. This increased severity of the functional deficit is evident in the degree of change between the weightbearing ratio and pad weightbearing, both of which at 5-, 7-, and 10-DPF are significantly higher for female mice with stabilized versus unstabilized fractures (**Figure 2B-C**). Accelerated functional recovery in female mice with stabilized fracture versus unstable fracture can also be observed for the pad weight-bearing on the fractured limb. Here the mice with stabilized fracture are no longer significantly different versus naïve control mice at 7– and 10-DPF, but are still significantly different from unstable fracture at the same timepoints (**Figure 2C**). Similarly, in the fractured hindlimb, for both step duration by 5-DPF (**Figure 2E**) and maximum paw speed by 10-DPF (**Figure 2F**), we recorded enhanced functional recovery in female mice with stabilized versus unstabilized.

In male mice, we observed largely similar changes in weight-bearing and kinematics after fracture. In males, we found that unstabilized fracture produces larger changes in weight-bearing than stabilized fracture at early fracture timepoints (**Figure 3B-C**). As in female mice, male mice with unstable fracture demonstrated significant changes in stepping correlation, with no changes observed after stabilized fracture (**Figure 3D**). Again, at early fracture timepoints, we observed significant increases in step duration and decreases in maximum paw speed (**Figure 3E-F**). For the kinematic parameters, one notable sex difference was that males lacked distinct changes in the step length, across fracture types (**Figure 3G**). In terms of the severity of the functional deficit and speed of functional recovery, males follow the similar pattern as the females above (**Figure 3A-G**).

**Figure 3.**
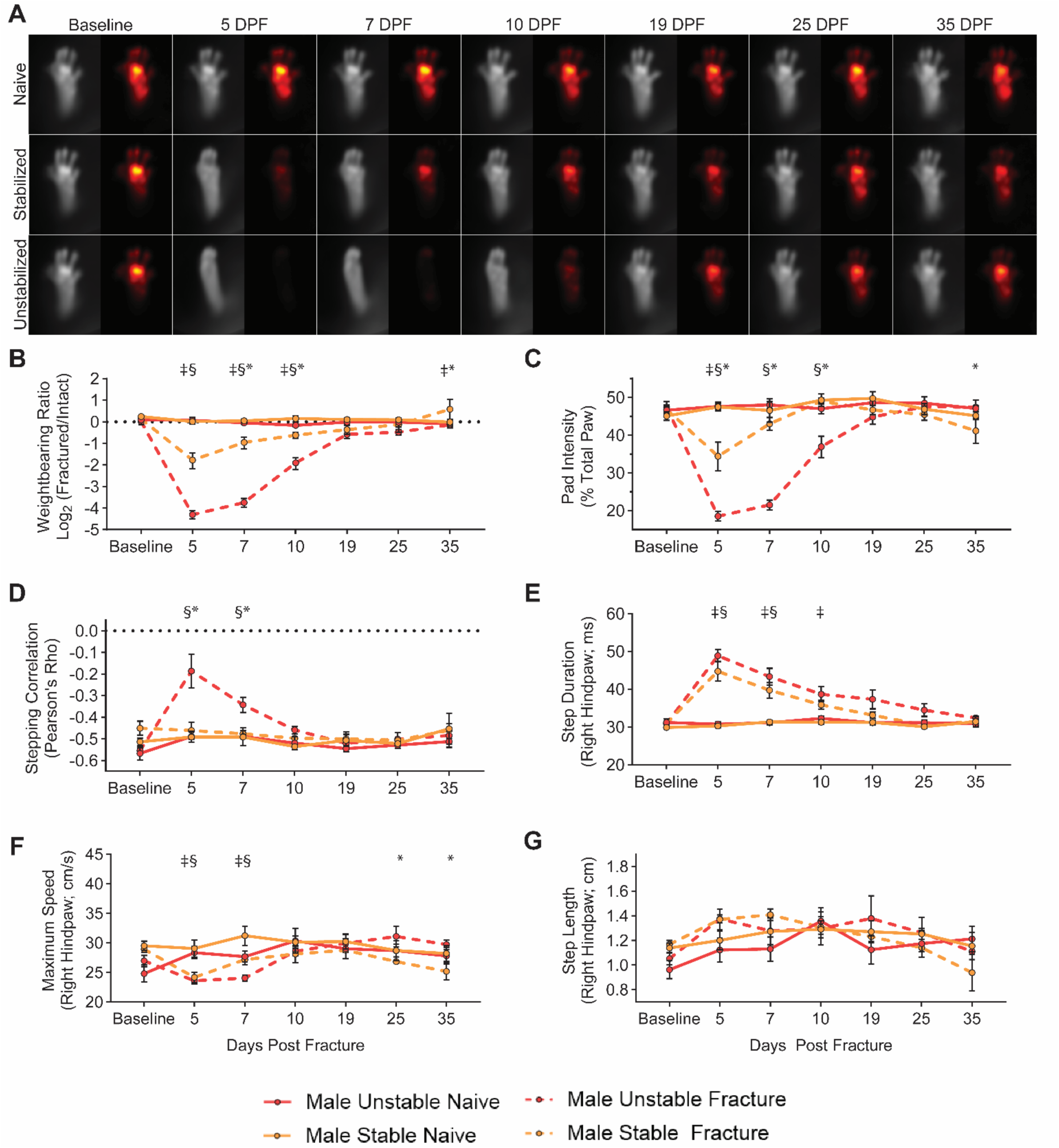
Differential functional impairments in male mice identified by fracture type. **(A)** Average images of paw placement (TL) and weightbearing (FTIR) of naïve, stabilized, and unstabilized fracture males at baseline and 5-, 7-, 10-, 19-, 25-, and 35-days post injury (DPI). Longitudinal analysis of weightbearing and kinematic parameters: **(B)** Weightbearing ratio of the fractured hindlimb to the intact hindlimb (Mixed-effects analysis, Fixed effects: Time (F(6, 42) = 10.79, p < 0.0001) Naïve vs. Fracture (F(1, 7) = 42.48, p = 0.0003). **(C)** Percentage of FTIR intensity localized to the pad of the hindpaw of the fractured hindlimb (Mixed-effects analysis, Fixed effects: Time (F(6, 42) = 24.90, p < 0.0001) Naïve vs. Fracture (F(1, 7) = 64.59, p < 0.0001). **(D)** Hindlimb stepping correlation (Mixed-effects analysis, Fixed effects: Time (F(6, 42) = 6.387, p < 0.0001) Naïve vs. Fracture (F(1, 7) = 11.30, p = 0.0121). **(E)** The average full-width, half-max duration of a single step, in milliseconds (Mixed-effects analysis, Fixed effects: Time (F(6, 42) = 33.92, p < 0.0001) Naïve vs. Fracture (F(1, 7) = 45.07, p = 0.0059). **(F)** Average maximum speed (cm/s) of right hindpaw during stepping (Mixed-effects analysis, Fixed effects: Time (F(6, 42) = 5.237, p = 0.0004) Naïve vs. Fracture (F(1, 7) = 2.898, p = 0.1325). **(G)** Average distance of each step of the right hindlimb in centimeters (Mixed-effects analysis, Fixed effects: Time (F(6, 42) = 5.508, p = 0.0003) Naïve vs. Fracture (F(1, 7) = 0.4581, p = 0.5203). Significant differences (p<0.05 after corrections for multiple comparisons) are indicated by: § – unstable fracture and respective naive control; ‡ – stable fracture and respective naive control; * – unstable and stable fracture.

### Male and Female Mice Exhibit Largely Similar Functional Changes After Fracture

We also sought to directly compare fracture-induced behavioral impairments and functional recovery patterns between female and male mice. Regarding weight-bearing metrics, we observed no significant sex-specific differences of the weight-bearing ratio or pad weight-bearing distribution during the main recovery period (5-25 DPF) (**Figure 4A-B**). A minor, yet significant, difference in stepping correlation was observed at 7 DPF identified between female and male mice with unstable fracture (**Figure 4C**). To account for observed differences in kinematic endpoints identified between the sexes of naïve animals (**Supplemental Figure 2**) and allow for direct comparisons not influenced by animal size and weight, we normalized kinematic endpoints (e.g., step length, step duration, maximum paw speed) in fractured mice to their respective naïve control groups for each sex and fracture type. Although there were no sex differences in the step duration or maximum speed (**Figure 4D-E**), we did find that at 5– and 19-DPF female mice have decreased normalized step length while this metric increased for males (**Figure 4F**). Taken together, these results demonstrate that, with a few minor exceptions, functional recovery from both stable and unstable fracture are largely similar in female and male mice.

**Figure 4.**
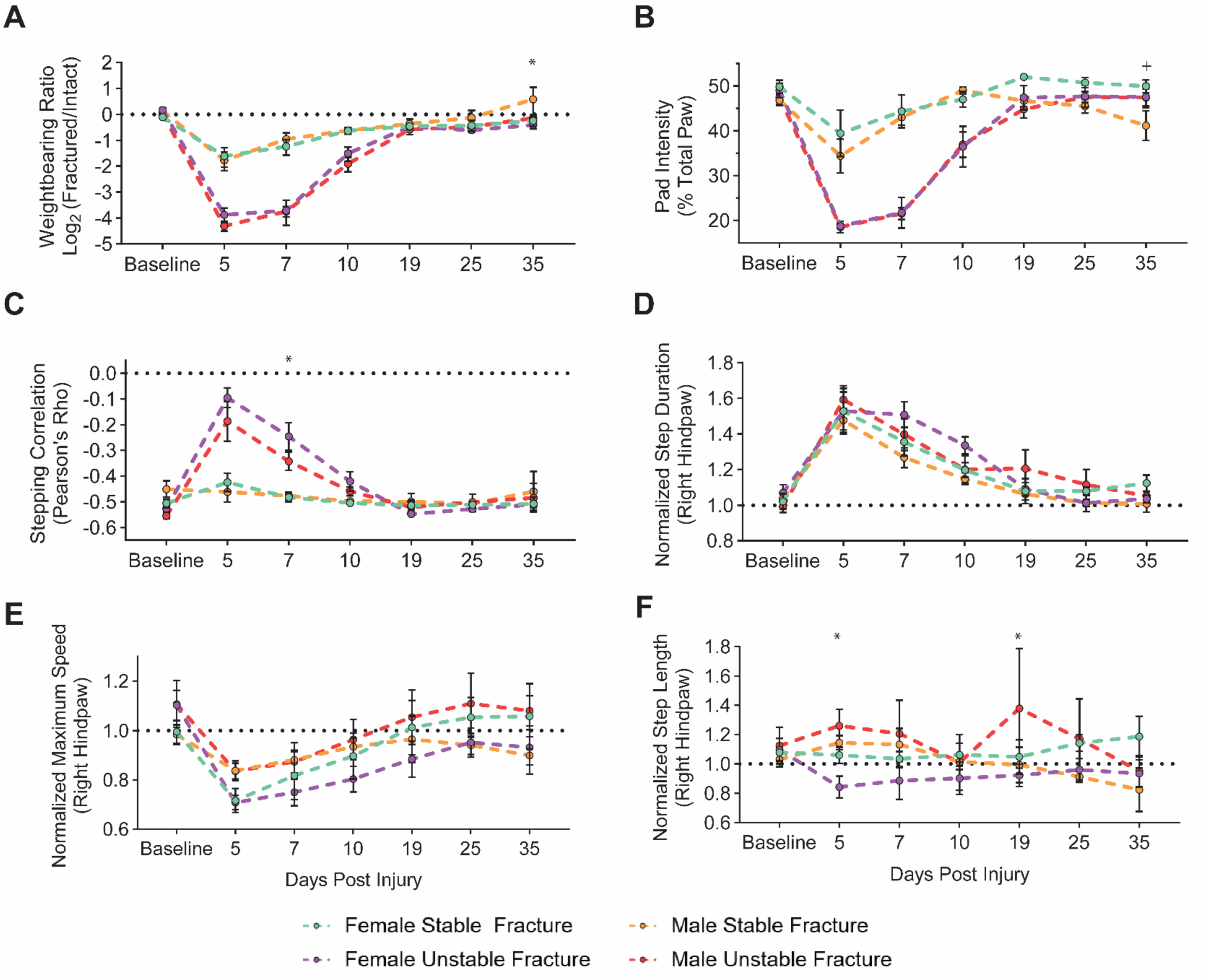
Male and female mice exhibit similar functional impairments and recovery within a fracture type. Longitudinal analysis of weightbearing and kinematic parameters comparing male and female mice after stable and unstable fracture: **(A)** Weightbearing ratio of the fractured hindlimb to the intact hindlimb (Mixed-effects analysis, Fixed effects: Time (F(6, 48) = 10.79, p < 0.0001) Stable vs. Unstable (F(1, 8) = 11.92, p = 0.2580). **(B)** Percentage of FTIR intensity localized to the pad of the hindpaw of the fractured hindlimb (Mixed-effects analysis, Fixed effects: Time (F(6, 48) = 60.35, p < 0.0001) Stable vs. Unstable (F(1, 8) = 16.65, p = 0.0035). **(C)** Hindlimb stepping correlation (Mixed-effects analysis, Fixed effects: Time (F(6, 48) = 26.38, p < 0.0001) Stable vs. Unstable (F(1, 8) = 14.9, p = 0.0055). **(D)** The normalized average full-width, half-max duration of a single step, in milliseconds (Mixed-effects analysis, Fixed effects: Time (F(6, 48) = 44.44, p < 0.0001) Stable vs. Unstable (F(1, 8) = 1.900, p = 0.2054). **(E)** Normalized average maximum speed (cm/s) of right hindpaw during stepping (Mixed-effects analysis, Fixed effects: Time (F(6, 48) = 15.30, p < 0.0001) Stable vs. Unstable (F(1, 8) = 0.3853, p = 0.8493). **(F)** Normalized average distance of each step of the right hindlimb in centimeters (Mixed-effects analysis, Fixed effects: Time (F(6, 48) = 0.7299, p < 0.6278) Stable vs. Unstable (F(1, 8) = 8.901e^-5^, p = 0.9927). Data in **D**, **E**, and **F** are all normalized to respective naïve control mice. Significant differences (p<0.05 after corrections for multiple comparisons) are indicated by: + – male and females with stable fracture; * – males and females with unstable fracture.

### Graph Theory-Based Behavioral Phenotyping Identifies Post-Fracture Recovery Window

Here, we sought to develop a unified, comprehensive metric that could integrate changes across all weightbearing and kinematic measurements produced by our analysis. As illustrated in the previous figures, individual metrics identify different time windows for functional recovery (for example, weight-bearing ratio vs. pad intensity). These metrics suggests that a unified metric could overcome these differences and provide an unbiased, global assessment of functional recovery during fracture healing. First, we used a z-scored heatmap to visualize how patterns of behaviors change across our entire dataset. Hierarchical clustering by behavioral metrics (x-axis) and fracture type, sex, and timepoint (y-axis) identified distinct groupings within our dataset, by fracture fixation and sex (**Supplemental Figure 4**). Similarly, using a pairwise correlation analysis to compare correlations across all the behavioral metrics between timepoints, we observed 3 distinct groupings within the data. These three groups are defined by male sex, female sex, and mice with fracture (**Supplemental Figure 5**).

We then used a graph theory framework to analyze and visualize changes across our combined dataset. Graph theory allows for a more comprehensive global representation of all possible combinations of pairwise relationships in a single analysis to identify patterns across the entire dataset, including behavioral state transitions associated with different datapoints within a group. In **Figure 5A**, each point on the graph is considered to be a “node”, and represents an individual recording (i.e., one recording from one mouse at one timepoint). Connections between nodes are called “edges” and signify the strength of the correlation between data (i.e., quantified behavioral recordings) from two nodes (i.e., animals). Clusters of nodes connected by a high density of edges indicate a higher degree of correlation, or similarity, between those individual behavioral patterns, and groupings of nodes that are farther from each other, connected by a lower density of edges, indicate less correlation. Using k-means clustering, we identified three separate clusters within our dataset (**Figure 5A**). The first two clusters correspond to non-fractured mice or late fracture timepoints (presumably when the animal has functionally recovered) for each sex (**cluster 1**: male, top left grouping of nodes *vs.* **cluster 2**: female, bottom left groupings of nodes). The third cluster is overwhelmingly represented by the early timepoints of the fractured mice (**cluster 3**: fracture, the right grouping of nodes). Importantly, as is shown in **Figure 5B**, nodes associated with individual animals shift their position within the graph over the time course of fracture related deficits and recovery. In early timepoints after the fracture, nodes are localized to the cluster defined by the fracture phenotype (cluster 3). However, as fracture healing occurs and mice functionally recover, nodes belonging to later timepoints progressively shift toward the naive groupings for each sex (**clusters 1 and 2**).

**Figure 5.**
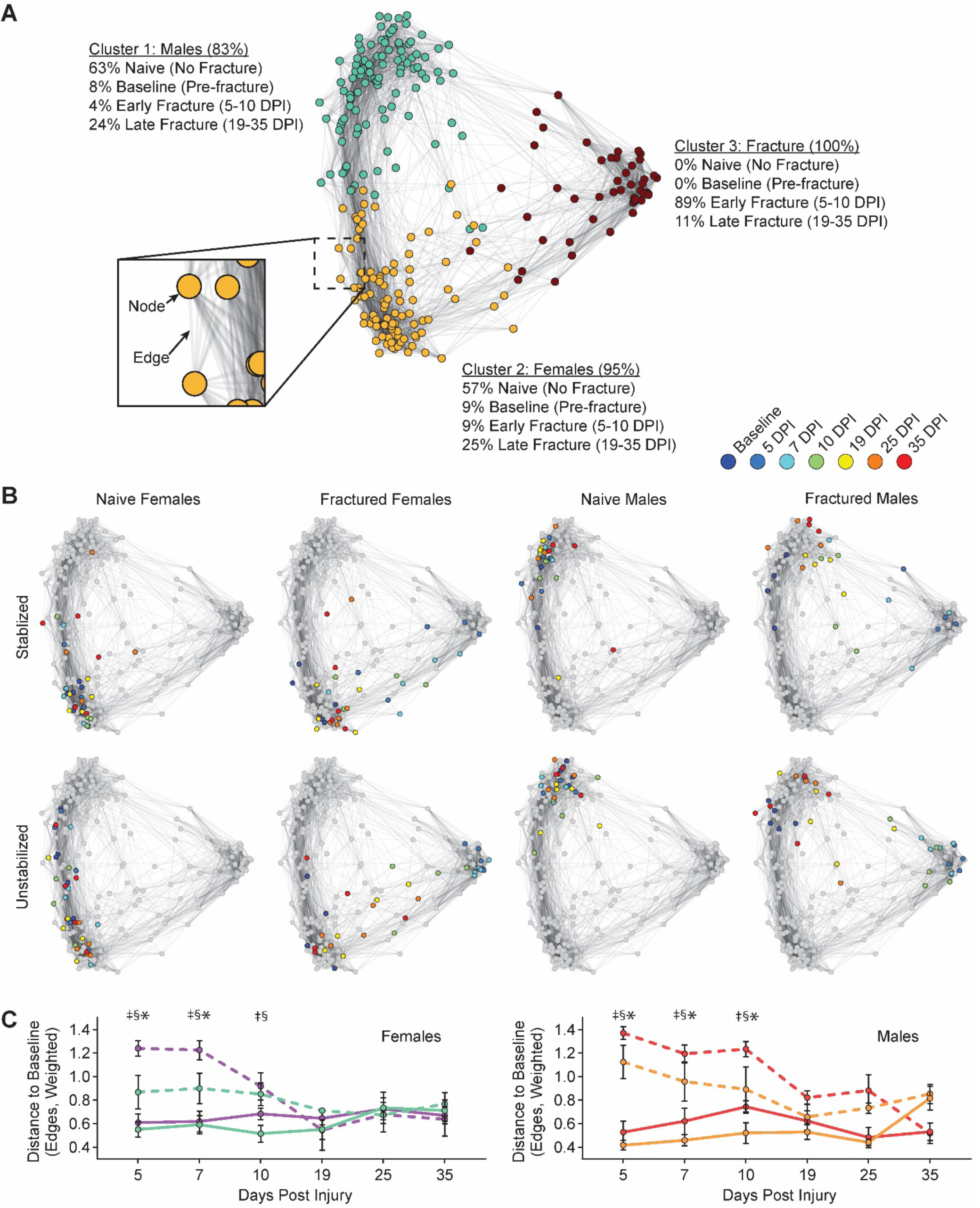
Graph Theory-Based Behavioral Phenotyping Identifies Post-Fracture Recovery Window. **(A)** Functional changes after tibia fracture are plotted using a graph theoretic approach. As represented in the cutout, individual recordings are represented as *nodes* in the graph, and positive correlations between sets of behaviors are represented as *edges*. k-means clustering identifies 3 clusters within the graph (nodes colored by cluster identity), with identities of nodes within each cluster indicated as percentages. **(B)** Representation of node identities on the graph, as indicated by sex and fracture conditions. Colored nodes indicated data from Baseline to 35 DPF for the indicated groupings (i.e., females with stable fracture), with all other nodes colored grey (i.e., those not in the indicated group). **(C)** For female (left) and male (right) mice, the degree of functional impairment is quantified as the distance of the post-fracture nodes to the baseline node (i.e., 5 DPI to baseline). Distance is measured in edges (weighted by the strength of the correlation between two nodes). Significant differences (p<0.05 after corrections for multiple comparisons) are indicated by: § – unstable fracture and respective naive control; ‡ – stable fracture and respective naive control; * – unstable and stable fracture.

To assess functional recovery, we quantified the dissimilarity of the behavioral states between any given timepoint (e.g., 5-DPF) and the baseline, unfractured state (**Figure 5C**). Here, we quantified the distance between nodes, which in graph theoretic terms refers to the shortest path length, in number of edges, that the node must traverse to reach the baseline node from the corresponding animal. We interpret the distance to the baseline node as how close the animal is to achieving functional recovery. We find that for both female and male mice, the distance to baseline for the fracture condition for both stable and unstable fractures is significantly different from naïve mice at early timepoints post fracture (5-10 DPF). However, at later timepoints (19-35 DPF), differences between fractured mice and naïve mice have largely dissipated.

## DISCUSSION

This study presents comprehensive behavioral phenotyping of the functional deficit and recovery time course of mice following fracture when fracture fixation and sex are varied in a controlled manner. To accomplish this, we combine longitudinal behavioral imaging analysis, machine learning, and graph theory analysis to rapidly identify and quantify variables that meaningfully translate to human changes in weightbearing and gait post-fracture. Consistent with clinical expectation, we found that tibia fractures with intramedullary stabilization present with less severe behavioral shifts compared to unstabilized fractures. On the other hand, we did not find that sex contributes to meaningful differences in functional healing.

To date, there has been a critical gap in technology that can quickly and reliably longitudinally quantify functional recovery in rodent models following fracture. Existing tools for preclinical gait analysis include DigiGait and TreadScan, which use transparent treadmills to identify abnormalities in rodent walking patterns, or the CatWalk, which requires that animals are trained to walk along a narrow glass platform and then produces pressure maps of the mouse paws.[35–37] The major limitation of these behavioral analyses is that the required training of each mouse is time consuming and also that the confined walking environment prevents animals from behaving naturally and ambulating freely. As our data clearly show, after a fracture, an animal’s balance and gait are compromised, limiting the effectiveness of forced gait tests due to their high dependence on highly stereotyped behaviors and parameters, including locomotor speed, consistent movement, and motivation.[33] Furthermore, data analysis from these methods is laborious, making it challenging to perform high throughput screening. Here, we present the first use of the Blackbox [31], a behavioral imaging system, to monitor weight-bearing and kinematic changes that occur during fracture injury and recovery. Our results clearly demonstrate that this novel behavioral phenotyping approach rapidly identifies and quantifies fracture-related behavioral changes, overcoming many of the challenges associated with other systems.

To validate our behavioral phenotyping methodology, we chose to use clinically meaningful variables expected to alter functional recovery following fracture. First, we changed the level of fracture stabilization. There is substantial evidence that motion across the healing fracture site plays a critical role in fracture repair. Limited motion promotes callus formation, but excessive motion can lead to non-union and chronic pain.[38–42] In clinical settings, the vast majority of tibia fractures are fixed surgically with intramedullary nails. This approach, in most cases, is believed to allow a sufficient degree of interfragmentary motion for optimal healing, and the nail allows patients to bear weight earlier.[43, 44] We model this scenario in the mice using intramedullary pin stabilization, which is well established in the field as a clinically relevant standard rodent model.[45] To model excessive motion we left the fractures unstabilized, which produces the same molecular healing response, but with a high degree of interfragmentary motion.[28] When comparing mice with an unstabilized fracture to a pin-stabilized fracture, as expected, we found distinct functional recovery patterns. Mice with unstabilized fractures demonstrated delayed weightbearing, take shorter steps, and their hind limb gait is more synchronous (not normal).

Due to the expected smaller degree of difference relative to fracture stabilization and relative paucity of data directly comparing female and male fracture recovery, we also evaluated sex-specific responses following fracture. Importantly, collectively, existing data remain inconclusive as to whether there are clinically meaningful sex-specific alterations in functional recovery or pain behavior following fracture. [46] We did not detect major sex-specific differences in functional recovery patterns. Consistent with our finding, Tawfik *et al* [47] did not find sex-specific differences in fracture responses using either von Frey fibers or gait analysis. The observed lack of behavioral differences between sexes was somewhat surprising as the Tawfik *et al* study and others have shown sex-specific divergence in innate and adaptive immune cell response after fracture, with a stronger immune response to injury in females.[47, 48] Clinical data have found sex differences in acute pain that are dependent upon the location of the fracture.[49] Of course, in non-fracture settings, there is evidence that females have higher incidence of pain, are more sensitive to painful stimulation as assessed in the laboratory, and are more likely to develop chronic pain (45% incidence versus 31%).[50–53]

Interpreting high dimensional datasets is an ongoing challenge, particularly when analyzing complex behavioral changes in response to injury. To address this challenge, we introduced a novel graph theoretic framework to analyze our combined datasets. This approach uniquely enables a global representation of large, complex datasets into a single framework that facilitates the estimation of simple and interpretable summary metrics. Our unified metric, the distance to the baseline node in graph theoretic space, made it possible to identify when fractured mice can reasonably be considered to have “recovered”. To the best of our knowledge, this is the first application of graph theory to analyze state changes in kinematic animal behaviors.

Despite minor, yet significant, differences in individual metrics at later fracture timepoints, our unified metric establishes that functional deficits persist at 10 DPF, but that both male and female mice are largely recovered by 19 DPF. This functional timeline coincides with the shift from cartilage to bone in the fracture callus.[7] The biological timeline of healing is well established in the fracture literature and it is known that at 5 DPF there is a robust pro-inflammatory response and a large hematoma within the fracture gap; as expected all mice at 5 DPF presented with severe functional deficits. Functional recovery begins between 7 and 10 DPF, correlating with the formation and maturation of cartilage within the fracture callus. By 19 DPF there is substantial trabecular bone formation bridging the fracture and we found that there are no longer substantial functional deficits in any of the mice. Other studies have shown that gait-related changes persist (for 4-6 weeks) in mice with a femur fracture, which heal more slowly than tibia fractures, however, these studies report the traditional single parameter view of functional recovery.[54, 55]

This study is a critical first step to addressing a technology gap in quantifying functional recovery following fracture. Our deep behavioral phenotyping was trained to rapidly and sensitively track interpretable, large data sets of kinematic and gait parameters. With graph theory we then create a unified metric of fracture recovery that is flexible and can easily be repurposed for other studies. In future studies, we plan to integrate our behavioral phenotyping approach with biological and/or mechanical healing parameters of the fracture. We also aim to expand our behavioral phenotyping approach so as to improve screening of therapeutics that can accelerate functional bone healing and analgesics that treat fracture pain.

## ACKNOWLEDGEMENTS

The authors thank Gina Baldoza for grant management and operations at the Orthopaedic Trauma Institute.

## FIGURE LEDGENDS

**Supplemental Figure 1.** Deeplabcut labeling strategy. Crosses correspond to points used for labeling for training pose estimation models within Deeplabcut.

**Supplemental Figure 2.** Sex differences in kinematic endpoints in naïve mice at baseline. Graphs illustrate sex differences in **(A)** body weight (g; student’s t-test, p<0.0001); **(B)** paw FTIR intensity (AU; student’s t-test, p=0.0092); **(C)** step duration (ms; student’s t-test, p<0.0001); **(D)** step length (cm; student’s t-test, p<0.0001); and **(E)** maximum paw speed (cm/s; student’s t-test, p<0.0001).

**Supplemental Figure 3.** Paw FTIR intensity correlates with mouse weight. Paw FTIR intensity significantly correlates with mouse weight during **(A)** stepping (Y =2065*X+21041; p=0.0147; R^2^=0.2082) and while **(B)** at rest, not stepping (Y=3730*X+398.3; p=0.0048; R^2^=0.2769). Solid line indicates best fit line after simple linear regression, and dotted lines indicate 95% confidence intervals.

**Supplemental Figure 4.** Z-scored heat map of all behavioral metrics over time. The x-axis of heatmap are represented by the different behavioral metrics assessed. The y-axis corresponds to data for a single condition (time, sex, fracture) averaged across all animals within that particular condition (M = male; F = female; BL = baseline; No-SFx or No-UFx indicate naïve controls; SFx and UFx indicate stabilized and unstablized fractures). In both dimensions, data are organized after ordering by hierarchical clustering analysis. Associated dendrograms are appended to each axis.

**Supplemental Figure 5.** Pairwise correlation matrix by behaviors over time. The x– and y-axis correspond to pairwise correlations across behavioral metrics between individual time points. Individual pixels are colored by correlation strength (Pearson’s correlation). Legend: (M = male; F = female; BL = baseline; No-SFx or No-UFx indicate naïve controls; SFx and UFx indicate stabilized and unstablized fractures). In both dimensions, data are organized after ordering by hierarchical clustering analysis, employing the same organization as the y-axis in **Supplemental Figure 4**.

