## Supplementary figures and images for "Deep behavioral phenotyping tracks functional recovery following tibia fracture in mice"

### Supplemental Figure 1

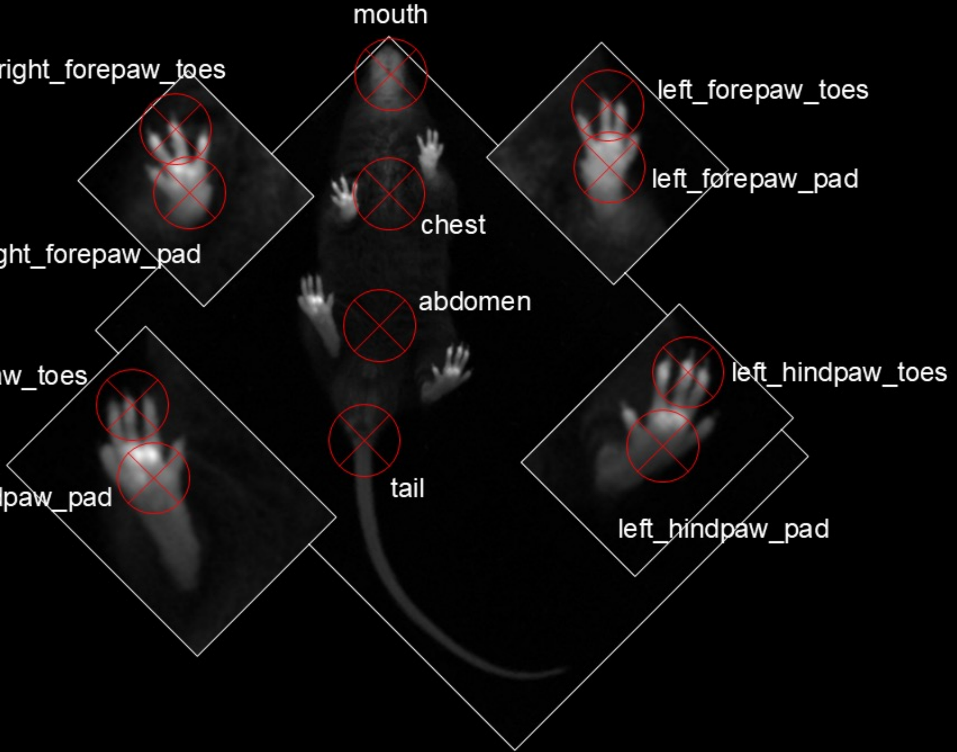

### Supplemental Figure 2

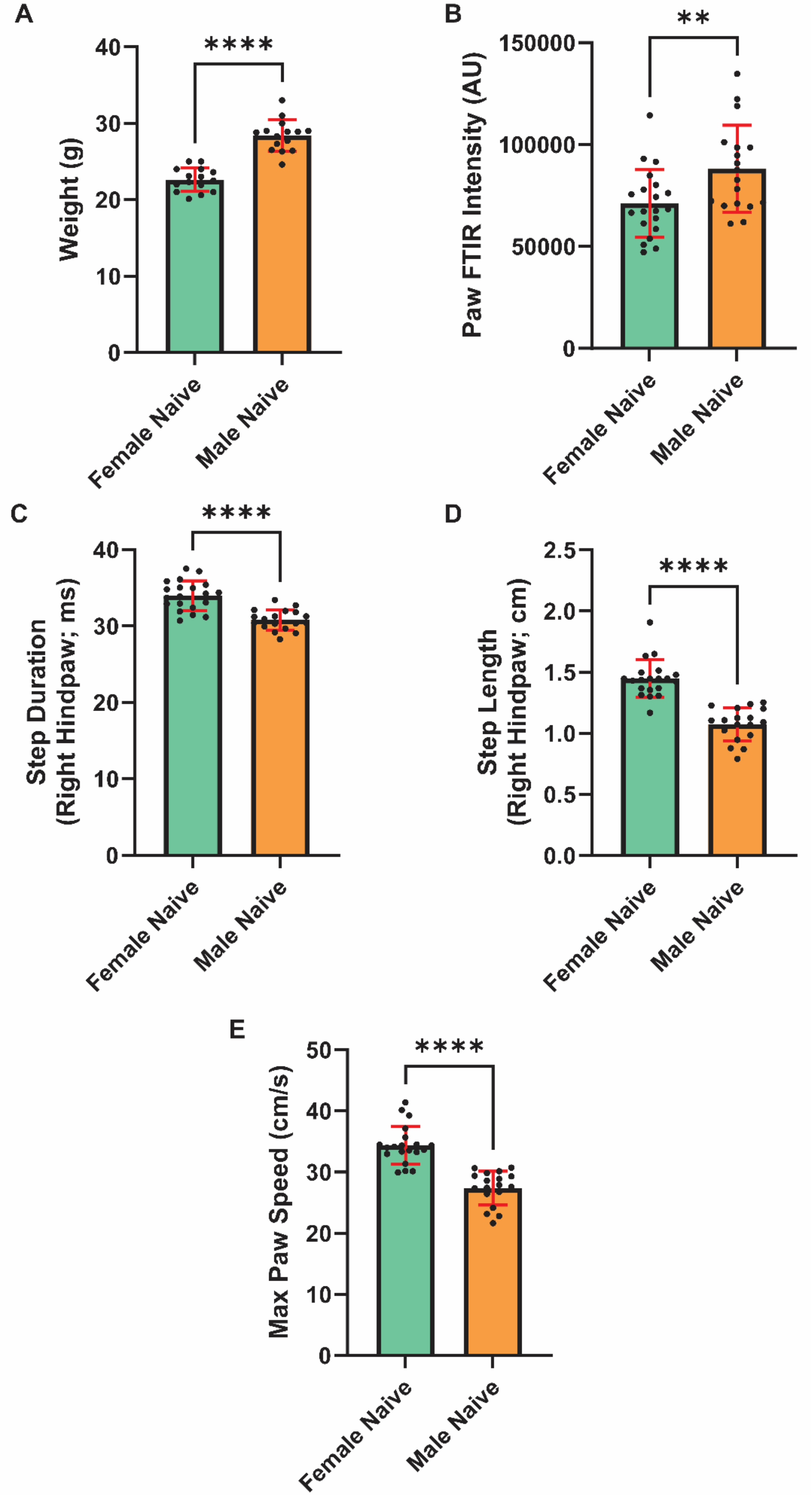

### Supplemental Figure 3

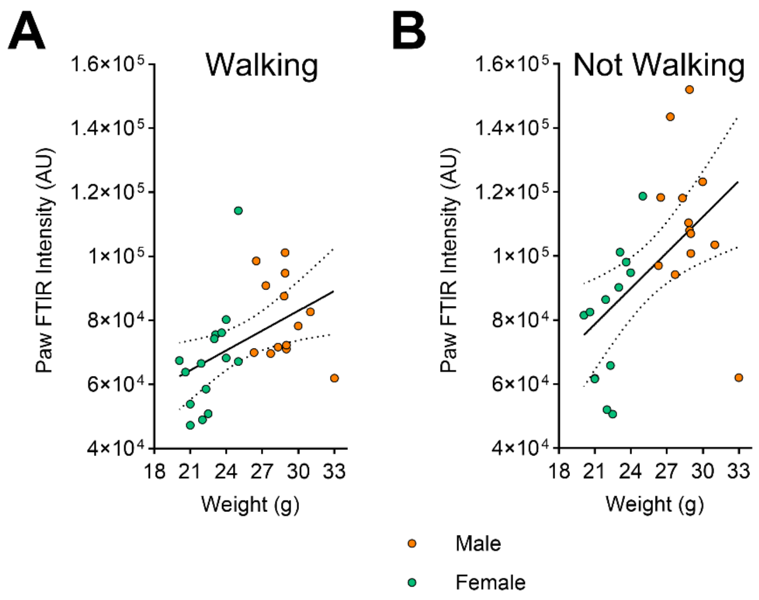

### Supplemental Figure 4

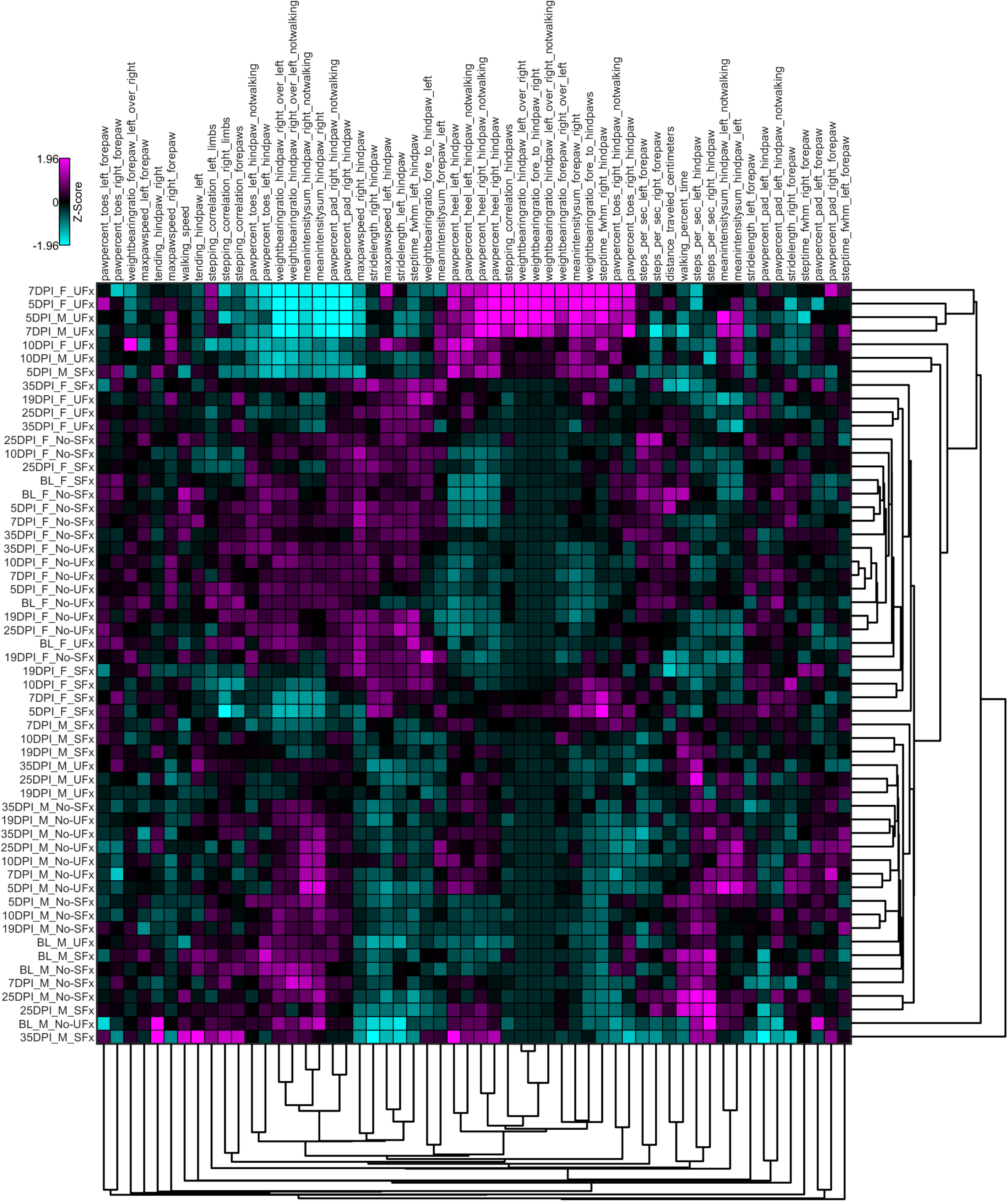

### Supplemental Figure 5

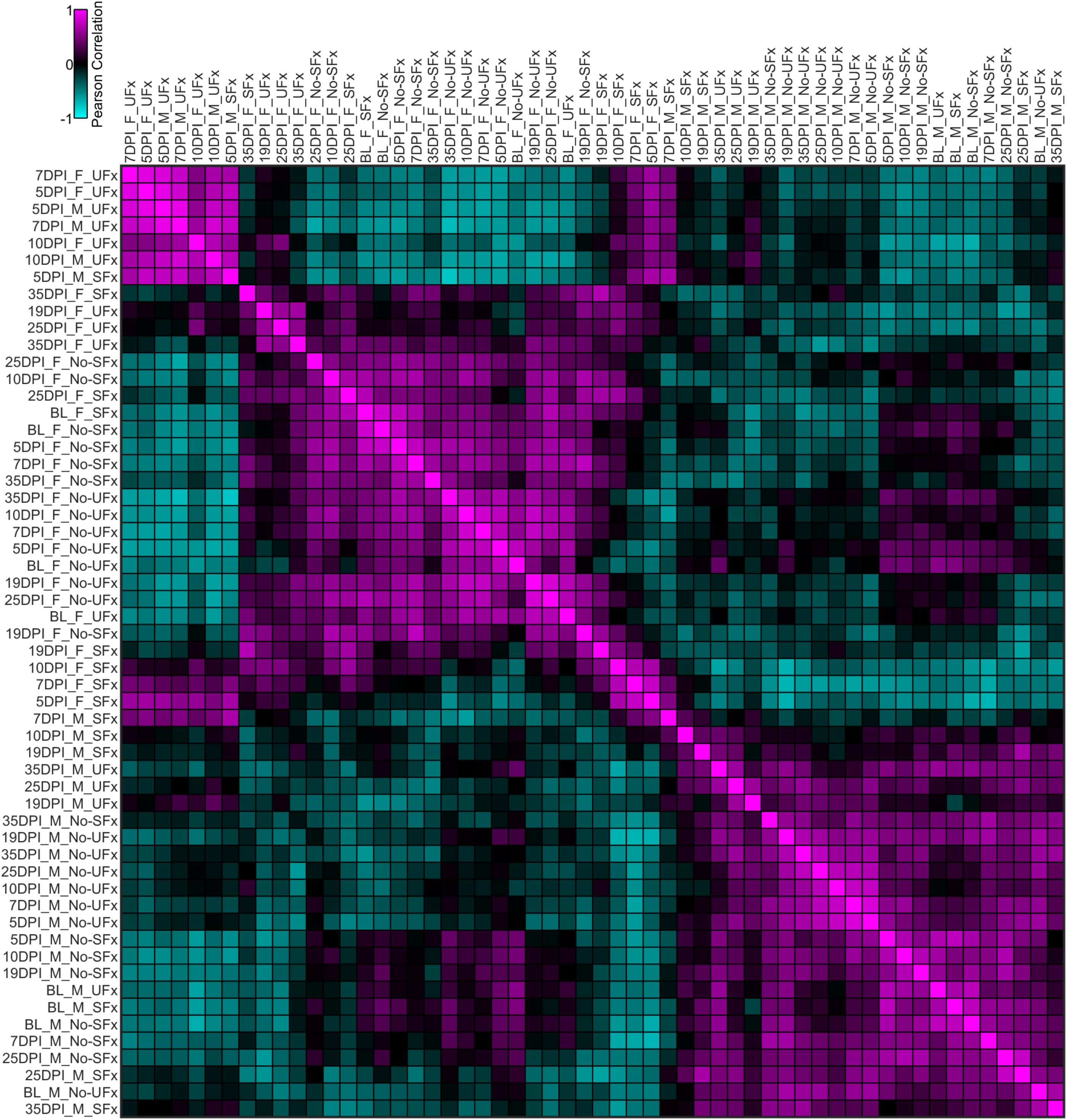
